# Drug resistance prediction for *Mycobacterium tuberculosis* with reference graphs

**DOI:** 10.1101/2023.05.04.539481

**Authors:** Michael B. Hall, Leandro Lima, Lachlan J. M. Coin, Zamin Iqbal

## Abstract

2.

The dominant paradigm for analysing genetic variation relies on a central idea: all genomes in a species can be described as minor differences from a single reference genome. However, this approach can be problematic or inadequate for bacteria, where there can be significant sequence divergence within a species.

Reference graphs are an emerging solution to the reference bias issues implicit in the “single-reference” model. Such a graph represents variation at multiple scales within a population – e.g., nucleotide- and locus-level.

The genetic causes of drug resistance in bacteria have proven comparatively easy to decode compared with studies of human diseases. For example, it is possible to predict resistance to numerous anti-tuberculosis drugs by simply testing for the presence of a list of single nucleotide polymorphisms and insertion/deletions, commonly referred to as a catalogue.

We developed DrPRG (Drug resistance Prediction with Reference Graphs) using the bacterial reference graph method Pandora. First, we outline the construction of a *Mycobacterium tuberculosis* drug resistance reference graph, a process that can be replicated for other species. The graph is built from a global dataset of isolates with varying drug susceptibility profiles, thus capturing common and rare resistance- and susceptible-associated haplotypes. We benchmark DrPRG against the existing graph-based tool Mykrobe and the haplotype-based approach of TBProfiler using 44,709 and 138 publicly available Illumina and Nanopore samples with associated phenotypes. We find DrPRG has significantly improved sensitivity and specificity for some drugs compared to these tools, with no significant decreases. It uses significantly less computational memory than both tools, and provides significantly faster runtimes, except when runtime is compared to Mykrobe on Nanopore data.

We discover and discuss novel insights into resistance-conferring variation for *M. tuberculosis* - including deletion of genes *katG* and *pncA* – and suggest mutations that may warrant reclassification as associated with resistance.

**Impact statement:** *Mycobacterium tuberculosis* is the bacterium responsible for tuberculosis (TB). TB is one of the leading causes of death worldwide; before the coronavirus pandemic it was the leading cause of death from a single pathogen. Drug-resistant TB incidence has recently increased, making the detection of resistance even more vital. In this study, we develop a new software tool to predict drug resistance from whole-genome sequence data of the pathogen using new reference graph models to represent a reference genome. We evaluate it on *M. tuberculosis* against existing tools for resistance prediction and show improved performance. Using our method, we discover new resistance-associated variations and discuss reclassification of a selection of existing mutations. As such, this work contributes to TB drug resistance diagnostic efforts. In addition, the method could be applied to any bacterial species, so is of interest to anyone working on antimicrobial resistance.

**Data summary:** **The authors confirm all supporting data, code and protocols have been provided within the article or through supplementary data files**.

The software method presented in this work, DrPRG, is freely available from GitHub under an MIT license at https://github.com/mbhall88/drprg. We used commit 9492f25 for all results via a Singularity[1] container from the URI docker://quay.io/mbhall88/drprg:9492f25.

All code used to generate results for this study are available on GitHub at https://github.com/mbhall88/drprg-paper. All data used in this work are freely available from the SRA/ENA/DRA and a copy of the datasheet with all associated phenotype information can be downloaded from the archived repository at https://doi.org/10.5281/zenodo.7819984 or found in the previously mentioned GitHub repository.

The *Mycobacterium tuberculosis* index used in this work is available to download through DrPRG via the command drprg index --download mtb@20230308 or from GitHub at https://github.com/mbhall88/drprg-index.

## 5. Introduction

Human industrialisation of antibiotic production and use over the last 100 years has led to a global rise in prevalence of antibiotic resistant bacterial strains. The phenomenon was even observed within patients in the first clinical trial of streptomycin as a drug for tuberculosis (TB) in 1948[2], and indeed as every new drug class has been introduced, so has resistance followed. Resistance mechanisms are varied, and can be caused by point mutations at key loci (e.g., binding sites of drugs[3,4]), frame-shifts rendering a gene non-functional[5], horizontal acquisition of new functionality via a new gene[6], or by up-regulation of efflux pumps to reduce the drug concentration within the cell[7].

Phenotypic and genotypic methods for detecting reduced susceptibility to drugs play complementary roles in clinical microbiology. Carefully defined phenotypic assays are used to give (semi)quantitative or binary measures of drug susceptibility; these have the benefit of being experimental, quantitative measurements, and are able to detect resistance caused by hitherto unknown mechanisms. Prediction of drug resistance from genomic data has different advantages. Detection of a single nucleotide polymorphism (SNP) is arguably more consistent than a phenotypic assay, as it is not affected by whether the resistance it causes is near some threshold defining a resistant/susceptible boundary. Additionally, combining sequence datasets from different labs is more reliable than combining different phenotypic datasets, and using sequence data allows one to detect informative genetic changes (e.g., a tandem expansion of a single gene to form an array, thus increasing dosage). More subtly, defining the cut-off to separate resistant from susceptible is only simple when the minimum inhibitory concentration distribution is a simple bimodal distribution; in reality it is sometimes a convolution of multiple distributions caused by different mutations, and genetic data is sometimes needed to deconvolve the data and choose a threshold[8,9].

The key requirement for a genomic predictor is to have an encodable understanding of the genotype-to-phenotype map. Research has focussed on clinically important pathogens, primarily *Escherichia coli, Klebsiella pneumoniae, Salmonella enterica, Pseudomonas aeruginosa* and *Mycobacterium tuberculosis* (MTB). The challenges differ across species; almost all bacterial species are extremely diverse, with non-trivial pan-genomes and considerable horizontal gene transfer causing transmission of resistance genes[10]. In these cases, species are so diverse that detection of chromosomal SNPs is affected heavily by reference bias[11]. Furthermore, there is an appreciable proportion of resistance which is not currently explainable through known SNPs or genes [12–14]. At the other extreme, MTB has almost no accessory genome, and no recombination or plasmids[15]. Resistance appears to be caused entirely by mutations, indels, and rare structural variants, and simple sets of rules (“if any of these mutations are present, or any of these genes inactivated, the sample is resistant”) work well for most drugs[16]. MTB has an exceptionally slow growth rate, meaning culture-based drug susceptibility testing (DST) is slow (2-4 weeks depending on media), and therefore sequencing is faster[17]. As part of the end TB strategy, the WHO strives towards universal access to DST[18], defining Target Product Profiles for molecular diagnostics[19,20] and publishing a catalogue of high-confidence resistance mutations intended to provide a basis for commercial diagnostics and future research[16]. There was a strong community-wide desire to integrate this catalogue into software for genotypic resistance prediction, although independent benchmarking confirmed there was still need for improvement[12]. Hence, there is a continuing need to improve the understanding of the genetic basis of resistance and integrate it into software for genotypic DST.

In this paper we develop and evaluate a new software tool for genotypic DST for MTB, built on a generic framework that can be used for any bacteria. Several tools have been developed previously[21–25]. Of these, only Mykrobe and TBProfiler work on Illumina and Nanopore data, and both have been heavily evaluated previously[22,23,26,27] - so we benchmark against these. Mykrobe uses de Bruijn graphs to encode known resistance alleles and thereby achieves high accuracy even on indel calls with Nanopore data[27]. However it is unable to detect novel alleles in known resistance genes, nor to detect gene truncation or deletion, which would be desirable. TBProfiler is based on mapping and variant calling (by default using Freebayes[28]), and detects gene deletions using Delly[29].

Our new software, called DrPRG (Drug resistance Prediction with Reference Graphs), builds on newer pan-genome technology than Mykrobe[11] using an independent graph for each gene in the catalogue, which makes it easier to go back-and-forth between VCF and the graph. To build an index, it takes as input a catalogue of resistant variants (a simple 4-column TSV file), a file specifying expert rules (e.g. any missense variant between codons X and Y in gene Z causes resistance to drug W), and a VCF of population variation in the genes of interest. This allows it to easily incorporate the current WHO-endorsed catalogue[16], which is conservative, and for the user to update the catalogue or rules with minimal effort. Finally, to provide resistance predictions, it takes a prebuilt index (an MTB one is currently provided) and sequencing reads (FASTQ).

We describe the DrPRG method, and to evaluate it, gather the largest MTB dataset of sequencing data with associated phenotype information and reveal novel insights into resistance-determining mutations for this species.

## 6. Methods

DrPRG is a command-line software tool implemented in the Rust programming language. There are two main subcommands: `build` for building a reference graph and associated index files, and `predict` for producing genotypic resistance predictions from sequencing reads and an index (from `build`).

### 6.1 Constructing a resistance-specific reference graph and index

The `build` subcommand of DrPRG requires a Variant Call Format (VCF) file of variants from which to build a reference graph, a catalogue of mutations that confer resistance or susceptibility for one or more drugs, and an annotation (GFF) and FASTA file of the reference genome.

For this work, we used the reference and annotation for the MTB strain H37Rv (accession NC_000962.3) and the default mutation catalogue from Mykrobe (v0.12.1)[12,26].

To ensure the reference graph is not biased towards a particular lineage or susceptibility profile, we selected samples from a VCF of 15,211 global MTB samples[30]. We randomly chose 20 samples from each lineage 1 through 4, as well as 20 samples from all other lineages combined. In addition, we included 17 clinical samples representing MTB global diversity (lineages 1-6)[31,32] to give a total of 117 samples. In the development phase of DrPRG we also found it necessary to add some common mutations not present in these 177 samples; as such, we added 48 mutations to the global VCF (these mutations are listed in archived repository – see Data summary). We did not add all catalogue mutations as there is a saturation point for mutation addition to a reference graph, and beyond this point, performance begins to decay[33].

The `build` subcommand turns this VCF into a reference graph by extracting a consensus sequence for each gene and sample. We use just those genes that occur in the mutation catalogue and include 100 bases flanking the gene. A multiple sequence alignment is constructed for each gene from these consensus sequences with MAFFT (v7.505)[34,35] and then a reference graph is constructed from these alignments with `make_prg` (v0.4.0)[11]. The final reference graph is then indexed with `pandora` [11].

### 6.2 Genotypic resistance prediction

Genotypic resistance prediction of a sample is performed by the predict subcommand of DrPRG. It takes an index produced by the `build` command (see Constructing a resistance-specific reference graph and index) and sequencing reads – Illumina or Nanopore are accepted. To generate predictions, DrPRG discovers novel variants (`pandora`), adds these to the reference graph (`make_prg` and MAFFT), and then genotypes the sample with respect to this updated graph (`pandora`). The genotyped VCF is filtered such that we ignore any variant with less than 3 reads supporting it and require a minimum of 1% read depth on each strand. Next, each variant is compared to the catalogue. If an alternate allele has been called that corresponds with a catalogue variant, resistance (‘R’) is noted for the drug(s) associated with that mutation. If a variant in the VCF matches a catalogue mutation, but the genotype is null (‘.’), we mark that mutation, and its associated drug(s), as failed (‘F’). Where an alternate allele call does not match a mutation in the catalogue, we produce an unknown (‘U’) prediction for the drug(s) that have a known resistance-conferring mutation in the relevant gene.

DrPRG also has the capacity to detect minor alleles and call minor resistance (‘r’) or minor unknown (‘u’) in such cases. Minor alleles are called when a variant (that has passed the above filtering) is genotyped as being the susceptible (reference) allele, but there is also read depth on the resistant (alternate) allele above a given minor allele frequency parameter (`--maf`; default is 0.1 for Illumina data). Minor allele calling is turned off by default for Nanopore data as we found it led to a drastic increase in the number of false positive calls (this is also the case for Mykrobe and TBProfiler).

When building the index for DrPRG and making predictions, we also accept a file of “expert rules” for calling variants of a certain class. A rule is associated with a gene, an optional position range, a variant type, and the drug(s) that rule confers resistance to. Currently supported variant types are missense, nonsense, frameshift, and gene absence.

The output of running predict is a VCF file of all variants in the graph and a JSON file of resistance predictions for each drug in the index, along with the mutation(s) supporting that prediction and a unique identifier to find that variant in the VCF file (see Supplementary Section S1 for an example). The reference graph gene presence/absence (as determined by `pandora`) is also listed in the JSON file.

### 6.3 Benchmark

We compare the performance of DrPRG against Mykrobe (v0.12.1)[26] and TBProfiler (v4.3.0)[22] for MTB drug resistance prediction. Mykrobe is effectively a predecessor of DrPRG; it uses genome graphs, in the form of de Bruijn graphs, to construct a graph of all mutations in a catalogue and then genotypes the reads against this graph. TBProfiler is a more traditional approach which aligns reads to a single reference genome and calls variants from that via aligned haplotype sequences.

A key part of such a benchmark is the catalogue of mutations, as this generally accounts for the majority of differences between tools[26]. As such, we use the same catalogue for all tools to ensure any differences are method-related - not catalogue disparities. The catalogue we chose is the default one provided in Mykrobe[12]. It is a combination of the catalogue described in Hunt *et al*. [26] and the category 1 and 2 mutation and expert rules from the 2021 WHO catalogue[16]. This catalogue contains mutations for 14 drugs: isoniazid, rifampicin, ethambutol, pyrazinamide, levofloxacin, moxifloxacin, ofloxacin, amikacin, capreomycin, kanamycin, streptomycin, ethionamide, linezolid, and delamanid.

We used Mykrobe and TBProfiler with default parameters, except for a parameter in each indicating the sequencing technology of the data as Illumina or Nanopore and the TBProfiler option to not trim data (as we do this in Quality control).

We compare the prediction performance of each program using sensitivity and specificity. To calculate these metrics, we consider a true positive (TP) and true negative (TN) as a case where a program calls resistance and susceptible, respectively, and the phenotype agrees; a false positive (FP) as a resistant call by a program but a susceptible phenotype, with false negatives (FN) being the inverse of FP. We only present results for drugs in the catalogue and where at least 10 samples had phenotypic data available.

To benchmark the runtime and memory usage of each tool, we used the Snakemake benchmark feature within our analysis pipeline[36].

### 6.4 Datasets

We gathered various MTB datasets where whole-genome sequencing data (Nanopore or Illumina) were available from public repositories (ENA/SRA/DRA) and associated phenotypes were accessible for at least one drug present in our catalogue[16,27,37–49]. All data was downloaded with `fastq-dl` (v1.1.1; https://github.com/rpetit3/fastq-dl).

### 6.5 Quality control

All downloaded Nanopore fastq files had adapters trimmed with `porechop` (v0.2.4; https://github.com/rrwick/Porechop), with the option to discard any reads with an adapter in the middle, and any reads with an average quality score below 7 were removed with `nanoq` (v0.9.0)[50]. Illumina reads were pre-processed with `fastp` (v0.23.2)[51] to remove adapter sequences, trim low quality bases from the ends of the reads, and remove duplicate reads and reads shorter than 30bp.

Sequencing reads were decontaminated as described in Hall *et al*.[27] and Walker *et al*.[16]. Briefly, sequenced reads were mapped to a database of common sputum contaminants and the MTB reference genome (H37Rv; accession NC_000962.3)[52] keeping only those reads where the best mapping was to H37Rv.

After quality control, we removed any sample with average read depth less than 15, or where more than 5% of the reads mapped to contaminants.

Lineage information was extracted from the TBProfiler results (see Benchmark).

### 6.6 Statistical Analysis

We used a Wilcoxon rank-sum paired data test from the Python library SciPy[53] to test for significant differences in runtime and memory usage between the three prediction tools.

The sensitivity and specificity confidence intervals were calculated with a Wilson’s score interval with a coverage probability of 95%.

## 7. Results

To benchmark DrPRG, Mykrobe, and TBProfiler, we gathered an Illumina dataset of 45,702 MTB samples with a phenotype for at least one drug. After quality control (see Quality control), this number reduced to 44,709. In addition, we gathered 142 Nanopore samples, of which 138 passed quality control. In Figure 1 we show all available drug phenotypes for those interested in the dataset, yet our catalogue does not offer predictions for all drugs listed (see Benchmark). Lineage counts for all samples that passed quality control and have a single, major lineage call can be found in Table 1.

**Figure 1:**
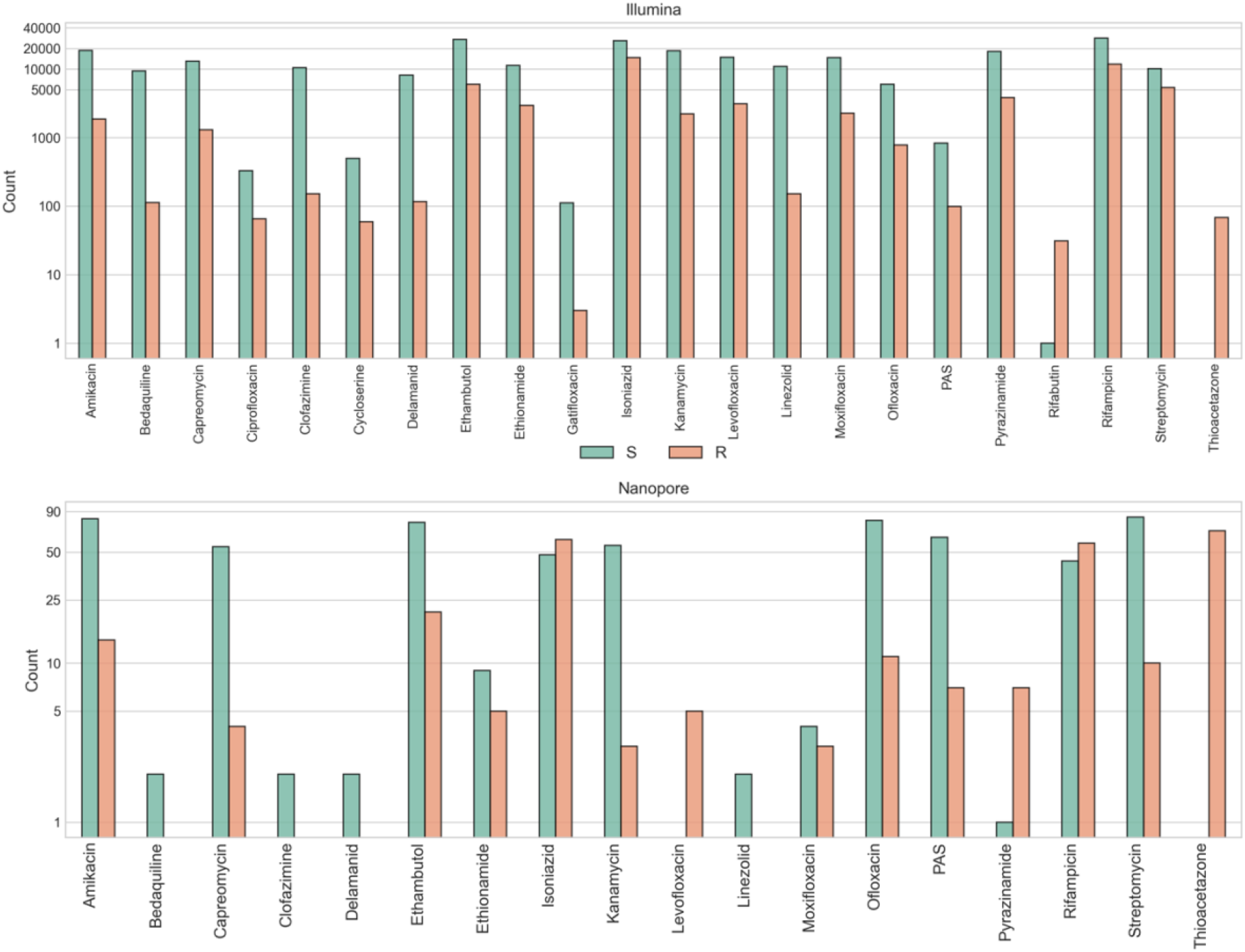
Drug phenotype counts for Illumina (upper) and Nanopore (lower) datasets. Bars are stratified and coloured by whether the phenotype is resistant (R; orange) or susceptible (S; green). Note, the y-axis is log-scaled. PAS=para-aminosalicylic acid

**Table 1:**
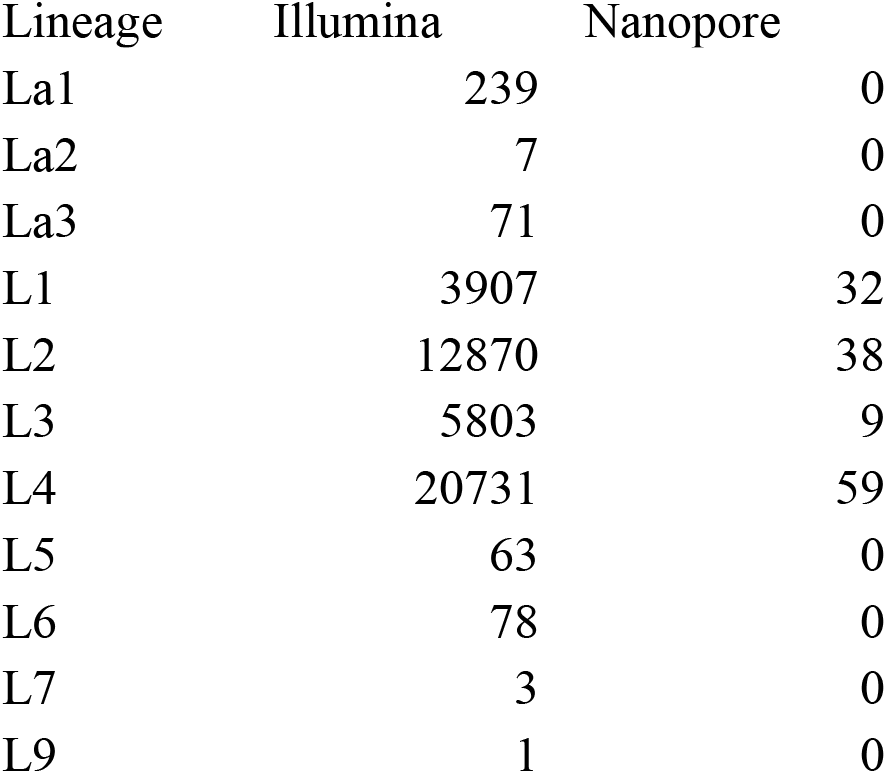
Lineage counts from the Illumina and Nanopore datasets, covering main lineages 1-9 (L1-L9) and the three livestock-associated lineages (La1-La3) as defined in [54].

| Lineage | Illumina | Nanopore |
| --- | --- | --- |
| La1 | 239 | 0 |
| La2 | 7 | 0 |
| La3 | 71 | 0 |
| L1 | 3907 | 32 |
| L2 | 12870 | 38 |
| L3 | 5803 | 9 |
| L4 | 20731 | 59 |
| L5 | 63 | 0 |
| L6 | 78 | 0 |
| L7 | 3 | 0 |
| L9 | 1 | 0 |

### 7.1 Sensitivity and specificity performance

We present the sensitivity and specificity results for Illumina data in Figure 2 and Suppl. Table S1 and the Nanopore data in Figure 3 and Suppl. Table S2.

**Figure 2:**
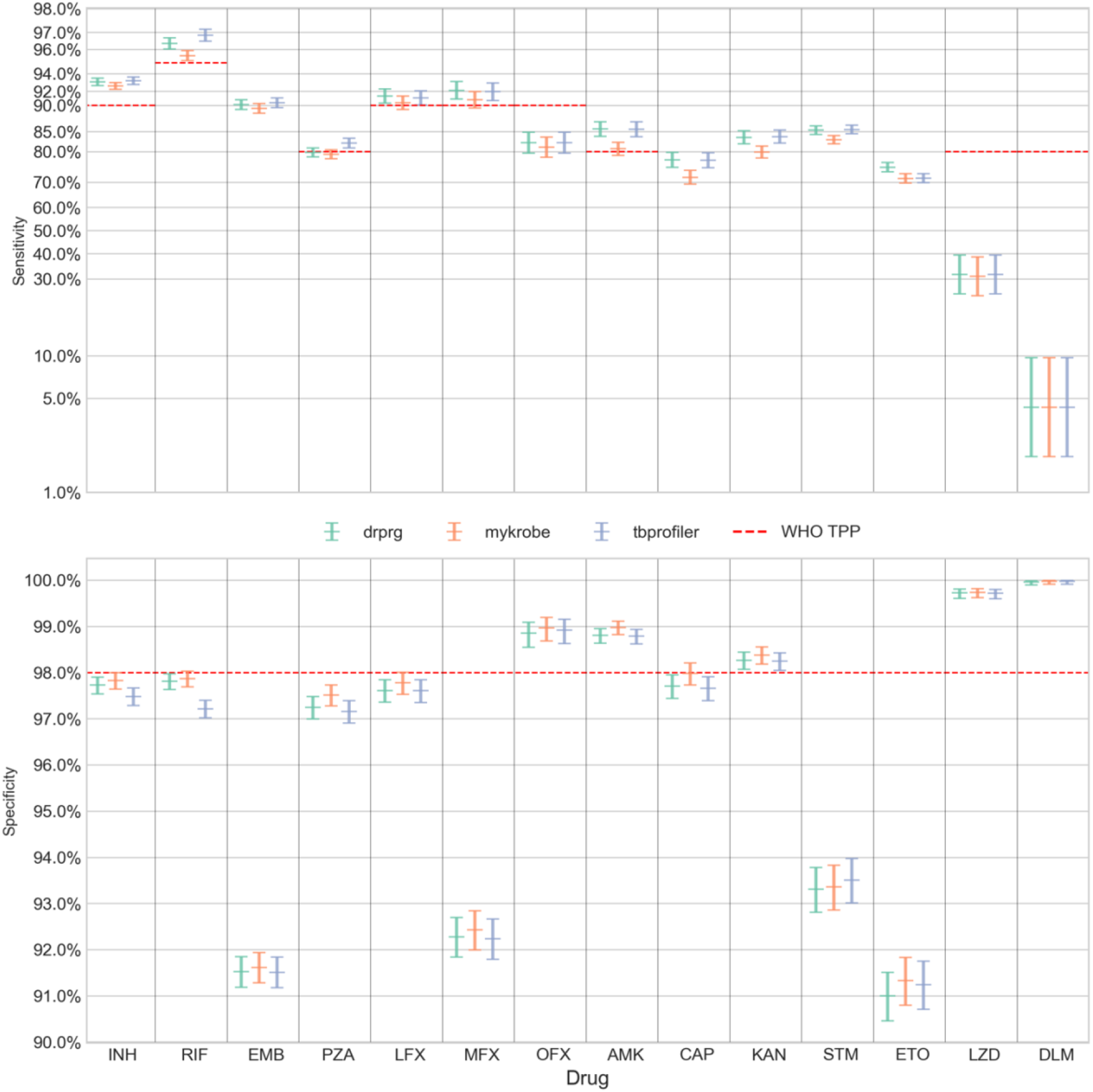
Sensitivity (upper panel; y-axis) and specificity (lower panel; y-axis) of resistance predictions for different drugs (x-axis) from Illumina data. Error bars are coloured by prediction tool. The central horizontal line in each error bar is the sensitivity/specificity and the error bars represent the 95% confidence interval. Note, the sensitivity panel’s y-axis is logit-scaled. This scale is similar to a log scale close to zero and to one (100%), and almost linear around 0.5 (50%). The red dashed line in each panel represents the minimal standard WHO target product profile (TPP; where available) for next-generation drug susceptibility testing for sensitivity and specificity. INH=isoniazid, RIF=rifampicin, EMB=ethambutol, PZA=pyrazinamide, LFX=levofloxacin, MFX=moxifloxacin, OFX=ofloxacin, AMK=amikacin, CAP=capreomycin, KAN=kanamycin, STM=streptomycin, ETO=ethionamide, LZD=linezolid, DLM=delamanid.

**Figure 3:**
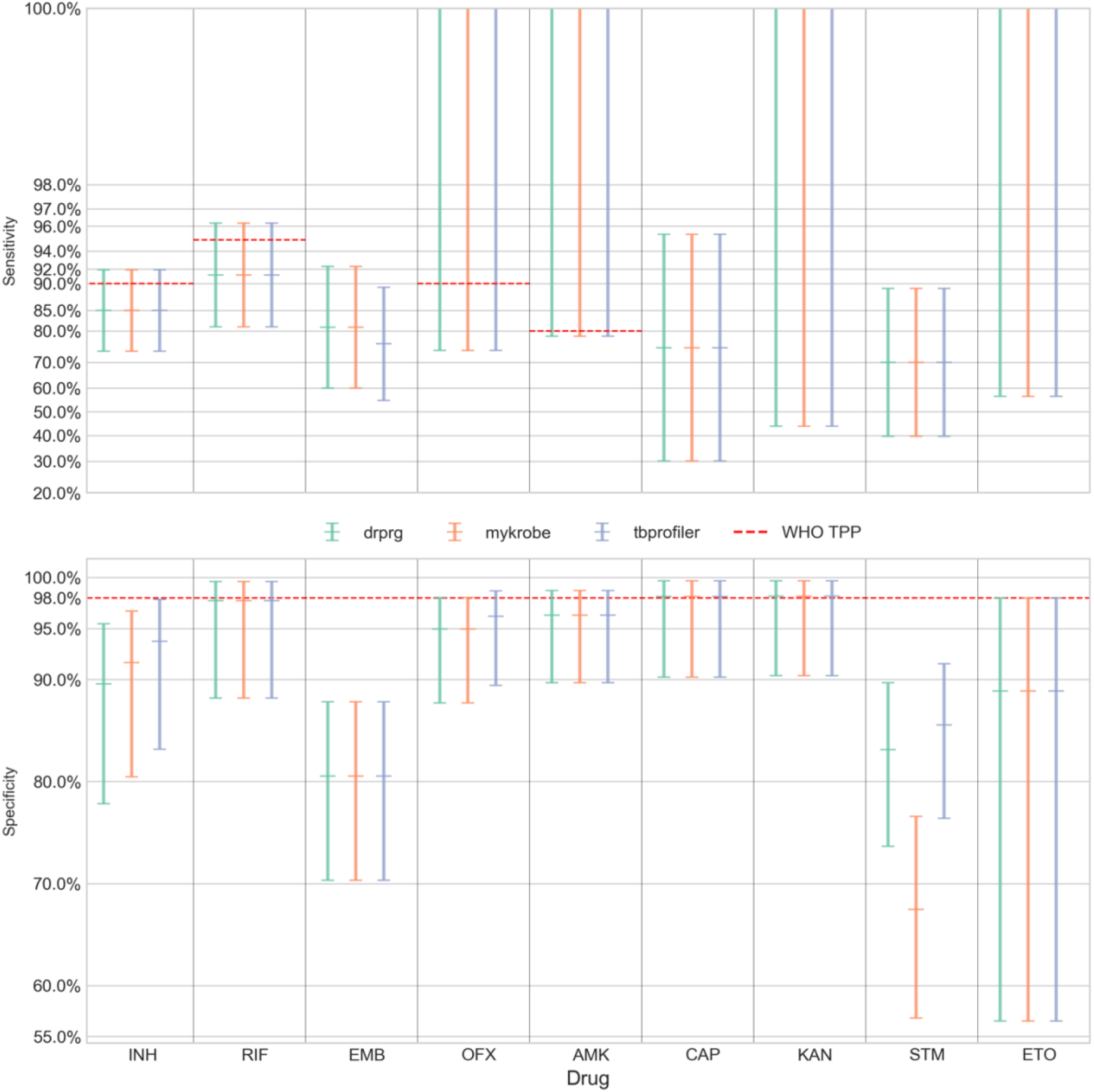
Sensitivity (upper panel; y-axis) and specificity (lower panel; y-axis) of resistance predictions for different drugs (x-axis) from Nanopore data. Error bars are coloured by prediction tool. The central horizontal line in each error bar is the sensitivity/specificity and the error bars represent the 95% confidence interval. Note, the sensitivity panel’s y-axis is logit-scaled. This scale is similar to a log scale close to zero and to one (100%), and almost linear around 0.5 (50%). The red dashed line in each panel represents the minimal standard WHO target product profile (TPP; where available) for next-generation drug susceptibility testing for sensitivity and specificity. INH=isoniazid, RIF=rifampicin, EMB=ethambutol, OFX=ofloxacin, AMK=amikacin, CAP=capreomycin, KAN=kanamycin, STM=streptomycin, ETO=ethionamide.

When comparing DrPRG’s performance to that of Mykrobe and TBProfiler, we look for instances where the confidence intervals do not overlap; indicating a significant difference. With Illumina data (Figure 2 and Suppl. Table S1), DrPRG achieves significantly greater sensitivity than Mykrobe for rifampicin (96.4% [96.0-96.7] vs. 95.6% [95.2-95.9]), streptomycin (85.3% [84.4-86.3] vs. 83.1% [82.1-84.1]), amikacin (85.6% [83.9-87.1] vs. 80.8% [78.9-82.5]), capreomycin (77.5% [75.2-79.7] vs. 71.8% [69.3-74.1]), kanamycin (83.7% [82.1-85.2] vs. 79.9% [78.2-81.5]), and ethionamide (75.2% [73.7-76.8] vs. 71.4% [69.7-73.0]), with no significant difference for all other drugs. In terms of sensitivity, there was no significant difference between DrPRG and TBProfiler except for ethionamide, where DrPRG was significantly more sensitive (75.2% [73.7-76.8] vs. 71.5% [69.8-73.1]). For specificity, there was no significant difference between the tools except that DrPRG and Mykrobe were significantly better than TBProfiler for rifampicin (97.8% [97.6-98.0] vs. 97.2% [97.0-97.4]). There was no significant difference in sensitivity or specificity for any drug with Nanopore data.

In both figures, we show the minimal requirements from the WHO target product profiles for sensitivity and specificity of genotypic drug susceptibility testing[19] as red dashed lines. Note, a sensitivity target is not specified by the WHO for ethambutol (EMB), capreomycin (CAP), kanamycin (KAN), streptomycin (STM), or ethionamide (ETO). For Illumina data, all tools’ predictions for rifampicin, isoniazid, levofloxacin, moxifloxacin and amikacin are above the sensitivity minimal requirement target. TBProfiler also exceeds the target for pyrazinamide, which DrPRG misses by 0.2%. No drug’s sensitivity target was achieved with Nanopore data. For specificity, the tools are all very similar and either exceed or fall below the threshold together (see Figure 2). The target of >98% is met by all tools on Illumina data only for ofloxacin, amikacin, linezolid, and delamanid. Mykrobe also exceeds the target for capreomycin. As such, amikacin is the only drug where both sensitivity and specificity performance exceed the minimal requirement of the WHO target product profiles. Only capreomycin and kanamycin specificity targets are exceeded (by all tools) with Nanopore data.

However, for Illumina data, we did find that likely sample-swaps or phenotype instability[55] could lead to some drugs being on the threshold of the WHO target product profiles. If we excluded samples where all three tools make a FP call for the strong isoniazid and rifampicin resistance-conferring mutations *katG* S315T (*n*=152) and *rpoB* S450L (*n*=119) [16] respectively, all three tools would exceed the isoniazid specificity target of 98% - thus meeting both sensitivity and specificity targets for isoniazid. In addition, DrPRG and Mykrobe would meet the rifampicin specificity target of 98% – leading to both targets being met for rifampicin for these two tools. As previously reported [55,56], we also found a lot of instability in the ethambutol result caused by *embB* mutations M306I (*n*=827) and M306V (*n*=519) being called for phenotypically susceptible samples (FP) by all three tools. Other frequent consensus FP calls included: *fabG1* c-15t, which is associated with ethionamide (*n*=441) and isoniazid (*n*=241) resistance; *rrs* a1401g, which is associated with resistance to capreomycin (*n*=241), amikacin (*n*=70), and kanamycin (*n*=48). In addition there were common false positives from *gyrA* mutations A90V and D94G, which are associated with resistance to the fluoroquinolones levofloxacin (*n*=108 and *n*=70, respectively), moxifloxacin (*n*=419 and *n*=349) and ofloxacin (*n*=19 and *n*=17), and are known to cause heteroresistance and minimum inhibitory concentrations (MIC) close to the critical concentration threshold[57–59].

### 7.2 Evaluation of potential additions to the WHO catalogue

False negatives are much harder to investigate as it is not known which mutation(s) were missed as they are presumably not in the catalogue if all tools failed to make a call. However, looking through those FNs where DrPRG makes an “unknown” resistance call, we note some potential mutations that may need reclassification or inclusion in the WHO catalogue. For delamanid FNs, we found five different nonsense mutations in the *ddn* gene in seven samples – W20^*^ (*n*=2), W27^*^ (*n*=1), Q58^*^ (*n*=1), W88^*^ (*n*=2), and W139^*^ (*n*=1) – none of which occurred in susceptible samples. We also found 13 pyrazinamide FN cases with a nonstop (stop-loss) mutation in *pncA* – this mutation type was also seen in two susceptible samples. Another *pncA* mutation, T100P, was also observed in 10 pyrazinamide FN samples and no susceptible samples. T100P only appears once in the WHO catalogue data (“solo” in a resistant sample). As such, it was given a grading of uncertain significance. As our dataset includes those samples in the WHO catalogue dataset, this means an additional nine isolates have been found with this mutation - indicating this may warrant an upgrade to ‘associated with resistance’. We found an interesting case of allele combinations, where nine ethambutol FN samples have the same two *embA* mutation c-12a and c-11a and *embB* mutation P397T - this combination is only seen in two susceptible samples. Interestingly, *embB* P397T and *embA* c-12a don’t appear in the WHO catalogue, but have been described as causing resistance previously[60]. Three *katG* mutations were also detected in isoniazid FN cases.

First, G279D occurs in eight missed resistance samples and no susceptible cases. This mutation is graded as ‘uncertain significance’ in the WHO catalogue and was seen solo in four resistant samples in that data. Singh *et al*. performed a protein structural analysis caused by this mutation and found it produced “an undesirable effect on the functionality of the protein”[61]. Second, G699E occurs in eight FN samples and no susceptible cases, but has a WHO grading of ‘uncertain significance’ based on six resistant isolates; thus, we add two extra samples to that count. And third, N138H occurs in 14 FN samples and one susceptible. In seven of these cases, it co-occurs with *ahpC* mutations t-75g (*n*=2) and t-76a (*n*=5). This mutation occurs in only three resistant isolates in the WHO catalogue dataset, giving it an uncertain significance, but we add a further 11 cases. This mutation has been found to cause a high isoniazid MIC and be associated with resistance[62,63].

### 7.3 Detection of large deletions

There are expert rules in the WHO catalogue which treat gene loss-of-function (any frameshift or nonsense mutation) in *katG, ethA, gid*, and *pncA* as causing resistance for isoniazid, ethionamide, streptomycin, and pyrazinamide, respectively[16]. Although examples of resistance caused by gene deletion are rare[64–68], with a dataset of this size (*n*=44,709), we can both evaluate these rules, and compare the detection power of DrPRG and TBProfiler for identifying gene deletions (Mykrobe does not, although in principle it could). In total we found 206 samples where DrPRG and/or TBProfiler identified deletions of *ethA, katG*, or *pncA*. Although many of these isolates did not have phenotype information for the associated drug (*n*=100), the results are nevertheless striking (Figure 4). Given the low false-positive rate of `pandora` for gene absence detection[11], these no-phenotype samples provide insight into how often gene deletions are occurring in clinical samples.

**Figure 4:**
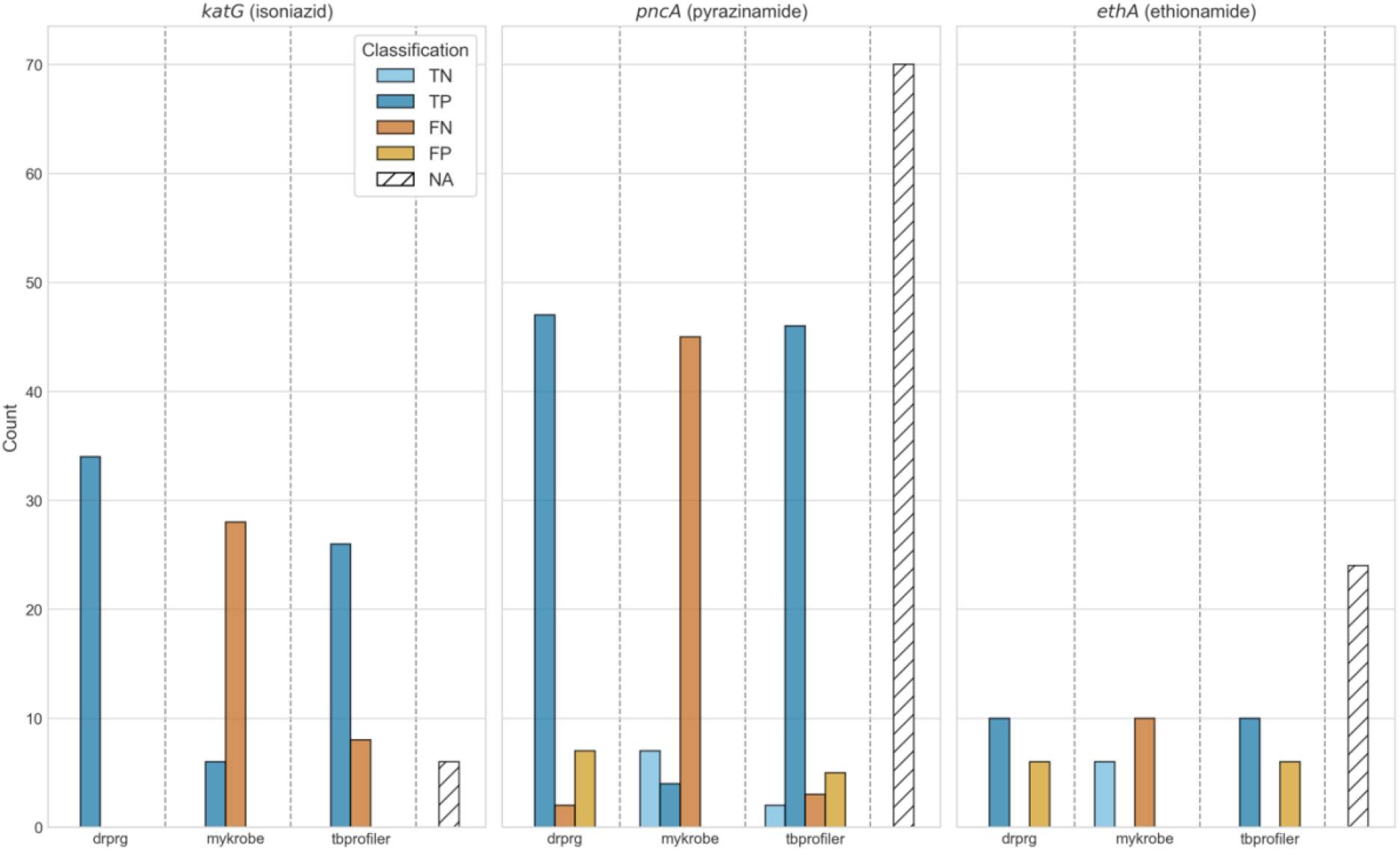
Impact of gene deletion on resistance classification. The title of each subplot indicates the gene and drug it effects. Bars are coloured by their classification and stratified by tool. Count (y-axis) indicates the number of gene deletions for that category. The NA bar (white with diagonal lines) indicates the number of samples with that gene deleted but no phenotype information for the respective drug. TP=true positive; FN=false negative; TN=true negative; FP=false positive; NA=no phenotype available.

Of the 34 isolates where *katG* was identified as being absent, and an isoniazid phenotype was available, all 34 were phenotypically resistant. DrPRG detected all 34 (100% sensitivity) and TBProfiler identified 26 (76.5% sensitivity). Deletions of *pncA* were detected in 56 isolates, of which 49 were phenotypically resistant. DrPRG detected 47 (95.9% sensitivity) and TBProfiler detected 46 (93.9% sensitivity). Lastly, *ethA* was found to be missing in 16 samples with an ethionamide phenotype, of which 10 were phenotypically resistant. Both DrPRG and TBProfiler correctly predicted all 10 (100% sensitivity). No *gid* deletions were discovered. We note that the TP calls made by Mykrobe were due to it detecting large deletions that are present in the catalogue, which is understandable given the whole gene is deleted.

We conclude that DrPRG is slightly more sensitive at detecting large deletions than TBProfiler (and Mykrobe) for *katG*, and equivalent for *pncA* and *ethA*. However we note that the WHO expert rule which predicts resistance to isolates missing specific genes appears more accurate for *katG* (100% of isolates missing the gene are resistant) than for *pncA* (87% resistant) and *ethA* (62.5% resistant).

### 7.4 Runtime and memory usage benchmark

The runtime and peak memory usage of each program was recorded for each sample and is presented in Figure 5. DrPRG (median 161 seconds) was significantly faster than both TBProfiler (307 seconds; *p*≤0.0001) and Mykrobe (230 seconds; *p*≤0.0001) on Illumina data. For Nanopore data, DrPRG (250 seconds) was significantly faster than TBProfiler (290 seconds; *p*≤0.0001), but significantly slower than Mykrobe (213 seconds; *p*=0.0347). In terms of peak memory usage, DrPRG (Illumina median peak memory 58MB; Nanopore 277MB) used significantly less memory than Mykrobe (1538MB; 1538MB) and TBProfiler (1463MB; 1990MB) on both Illumina and Nanopore data (*p*≤0.0001 for all comparisons).

**Figure 5:**
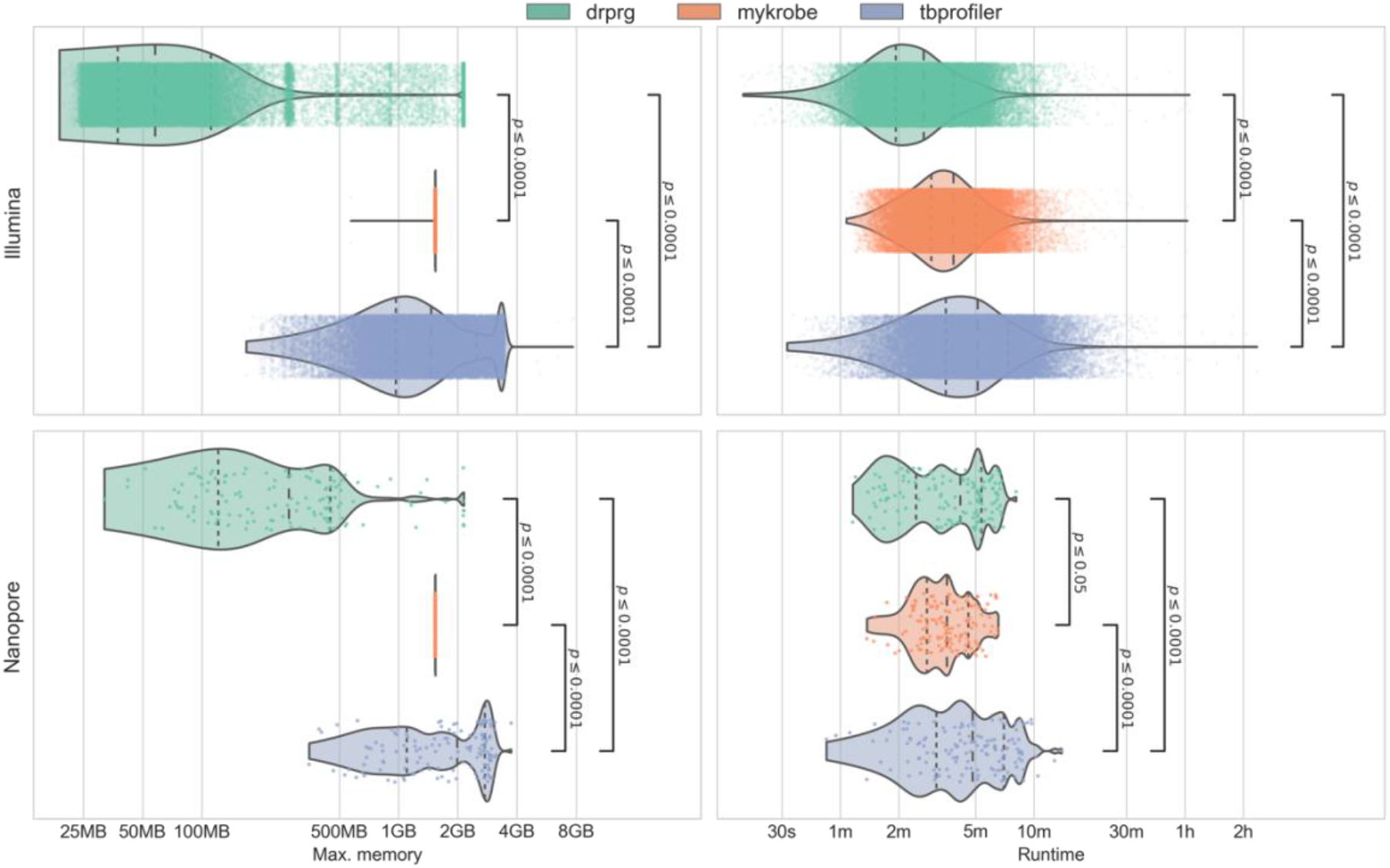
Benchmark of the maximum memory usage (left panels) and runtime (right panels) from Illumina (upper row) and Nanopore (lower row) data. Each point and violin is coloured by the tool, with each point representing a single sample. Statistical annotations are the result of a Wilcoxon rank-sum paired data test on each pair of tools. Dashed lines inside the violins represent the quartiles of the distribution. Note, the x-axis is log-scaled.

## 8. Discussion

In this work, we have presented a novel method for making drug resistance predictions with reference graphs. The method, DrPRG, requires only a reference genome and annotation, a catalogue of resistance-conferring mutations, a VCF of population variation from which to build a reference graph, and (optionally) a set of rules for types of variants in specific genes which cause resistance. We apply DrPRG to the pathogen *M. tuberculosis*, for which there is a great deal of information on the genotype/phenotype relationship, and a great need to provide good tools which implement and augment current and forthcoming versions of the WHO catalogue. We illustrate the performance of DrPRG against two existing methods for drug resistance prediction – Mykrobe and TBProfiler.

We benchmarked the methods on a high-quality Illumina sequencing dataset with associated phenotype profiles for 44,709 MTB genomes; the largest known dataset to-date[16]. All tools used the same catalogue and rules, and for most drugs, there was no significant difference between the tools. However, DrPRG did have a significantly higher specificity than TBProfiler for rifampicin predictions, and sensitivity for ethionamide predictions. DrPRG’s sensitivity was also significantly greater than Mykrobe’s for rifampicin, streptomycin, amikacin, capreomycin, kanamycin, and ethionamide. Evaluating detection of gene loss, we found DrPRG was more sensitive to *katG* deletions than TBProfiler.

We also benchmarked using 138 Nanopore-sequenced MTB samples with phenotype information, but found no significant difference between the tools. This Nanopore dataset was quite small and therefore the confidence intervals were large for all drugs. Increased Nanopore sequencing over time will provide better resolution of the overall sensitivity and specificity values and improve the methodological nuances of calling variants from this emerging, and continually changing, sequencing technology.

DrPRG also used significantly less memory than Mykrobe and TBProfiler on both Nanopore and Illumina data. In addition, the runtime of DrPRG was significant faster than both tools on Illumina data and faster than TBProfiler on Nanopore data. While the absolute values for memory and runtime for all tools mean they could all easily run on common computers found in the types of institutions likely to run them, the differences for the Nanopore data warrant noting. As Nanopore data can be generated “in the field”, computational resource usage is critical. For example, in a recent collaboration of ours with the National Tuberculosis program in Madagascar[27], Nanopore sequencing and analysis are regularly performed on a laptop, meaning memory usage is sometimes a limiting factor. DrPRG’s median peak memory was 277MB, meaning it can comfortably be run on any laptop and other mobile computing devices[69].

It is clear from the Illumina results that more work is needed to understand resistance-conferring mutations for delamanid and linezolid. However, we did find that nonsense mutations in the *ddn* gene appear likely to be resistance-conferring for delamanid – as has been noted previously[39,70–72]. We also found a novel (likely) mechanism of resistance to pyrazinamide - a nonstop mutation in *pncA*. Phenotype instability in *embB* at codon 306 was also found to be the main driver in poor ethambutol specificity, as has been noted elsewhere[55,56], indicating the need to further investigate cofactors that may influence the phenotype when mutations at this codon are present.

Gene absence/deletion detection allowed us to confirm that the absence of *katG* – a mechanism which is rare in clinical samples[64–67,73] - is highly likely to confer resistance to isoniazid. Additionally, we found that the absence of *pncA* is likely to cause resistance to pyrazinamide, as has been noted previously[68]. One finding that requires further investigation is the variability in ethionamide phenotype when *ethA* is absent. We found that only 63% of the samples with *ethA* missing, and an ethionamide phenotype, were resistant. An *et al*. have suggested that *ethA* deletion alone does not always cause resistance and there might be an alternate pathway via *mshA*[74].

Given the size of the Illumina dataset used in this work, the results provide a good marker of Illumina whole-genome sequencing’s ability to replace traditional phenotyping methods. With the catalogue used in this study, DrPRG meets the WHO’s target product profile for next-generation drug-susceptibility testing for both sensitivity and specificity for amikacin, and sensitivity only for rifampicin, isoniazid, levofloxacin, and moxifloxacin. However, if we exclude cases where all tools call *rpoB* S450L or *katG* S315T for phenotypically susceptible samples (these are strong markers of resistance[16] and therefore we suspect sample-swaps or phenotype error[75]), DrPRG also meets the specificity target product profile for rifampicin and isoniazid. For the other first-line drugs ethambutol and pyrazinamide, ethambutol does not have a WHO target and DrPRG’s sensitivity is 0.2% below the WHO target (although the confidence interval spans the target), while the specificity target is missed by 0.8%.

The primary limitation of the DrPRG method relates to minor allele calls. DrPRG uses `pandora` for novel variant discovery, which combines a graph of known population variants (which can be detected at low frequency) with *de novo* detection of other variants if present at above ∼50% frequency. Thus, it can miss minor allele calls if the allele is absent from its reference graph. While this issue did not impact most drugs, it did account for the majority of cases where DrPRG missed pyrazinamide-resistant calls (in *pncA*), but the other tools correctly called resistance. Unlike most other genes, where there are a relatively small number of resistance-conferring mutations, or they’re localised to a specific region (e.g. the rifampicin-resistance determining region in *rpoB*), resistance-conferring mutations are numerous - with most being rare - and distributed throughout *pncA*[16,76,77]. Adding all of these mutations will, and does, lead to decreased performance of the reference graph[33], and so improving minor allele calling for pyrazinamide remains a challenge we need to revisit in the future.

One final limitation is the small number of Nanopore-sequenced MTB isolates with phenotypic information. In order to get a clearer picture of the sensitivities and specificities this sequencing technology can provide, we need much larger and more diverse data.

In conclusion, DrPRG is a fast, memory frugal software program that can be applied to any bacterial species. We showed that on MTB, it performs as well as, or better than two other commonly used tools for resistance prediction. We also collected and curated the largest dataset of MTB Illumina-sequenced genomes with phenotype information and hope this will benefit future work to improved genotypic drug susceptibility testing for this species. While we applied DrPRG to MTB in this study, it is a framework that is agnostic to the species.

MTB is likely one of the bacterial species with the least to gain from reference graphs given its relatively conserved (closed) pan-genome compared to other common species[78]. As such, we expect the benefits and performance of DrPRG to improve as the openness of the species’ pan-genome increases[11]; especially given its good performance on a reasonably closed pan-genome.

## 9. Author statements

### 9.1 Author contributions

M.B.H: conceptualisation, data curation, formal analysis, investigation, methodology, resources, software, visualisation, writing – original draft, writing – review & editing. L.L: resources, software, writing – review & editing. L.J.M.C: funding acquisition, methodology, supervision, writing – review & editing. Z.I: conceptualisation, funding acquisition, methodology, supervision, writing – original draft, writing – review & editing.

## 9.4 Acknowledgements

We thank Kerri M. Malone for sharing her MTB knowledge through many discussions and critiquing the final manuscript. We also thank Martin Hunt and Jeff Knaggs for facilitating access to the CRyPTIC VCFs. Finally, we would like to acknowledge Timothy Walker for his clinically-relevant MTB advice.

