## Supplementary Material for "Drug resistance prediction for *Mycobacterium tuberculosis* with reference graphs"

### Contents

**S1 Example DrPRG prediction output** 2

**S2 Sensitivity and specificity performance** 4

### List of Tables

S1 Comparison of resistance predictions from Illumina data for three tools. FN=number of false negatives, R=number of resistant isolates, FP=number of false positives, S=number of susceptible isolates, CI=confidence interval. . . . . . . . . . . . . . . . . . . . . . 5

S2 Comparison of resistance predictions from Nanopore data for three programs. FN=number of false negatives, R=number of resistant isolates, FP=number of false positives, S=number of susceptible isolates, CI=confidence interval. . . . . . . . . . . . . 6

### S1 Example DrPRG prediction output

This is a trimmed (toy) example of the JSON output from DrPRG.


```
{
  "genes": {
    "absent": [
      "ahpC"
    ],
    "present": [
      "embA",
      "embB",
      "ethA",
      "fabG1",
      "gid",
      "gyrA",
      "gyrB",
      "inhA",
      "katG",
      "pncA",
      "rpoB",
      "rrs"
    ]
  },
  "sample": "toy",
  "susceptibility": {
    "Amikacin": {
      "evidence": [
        {
          "gene": "rrs",
          "residue": "DNA",
          "variant": "A1401X",
          "vcfid": "b815ed3f"
        }
      ],
      "predict": "F"
    },
    "Ethambutol": {
      "evidence": [
        {
          "gene": "embB",
          "residue": "PROT",
          "variant": "M306I",
          "vcfid": "a290b118"
        }
      ],
```

```
      "predict": "r"
    },
    "Ethionamide": {
      "evidence": [
        {
          "gene": "ethA",
          "residue": "PROT",
          "variant": "A381P",
          "vcfid": "169f75d4"
        }
      ],
      "predict": "U"
    },
    "Isoniazid": {
      "evidence": [
        {
          "gene": "fabG1",
          "residue": "DNA",
          "variant": "G-17T",
          "vcfid": "de9b689e"
        },
        {
          "gene": "katG",
          "residue": "PROT",
          "variant": "S315T",
          "vcfid": "acaa8ca2"
        }
      ],
      "predict": "R"
    },
    "Levofloxacin": {
      "evidence": [],
      "predict": "S"
    }
  },
  "version": {
    "drprg": "0.1.1",
    "index": "20230308"
  }
}
```


The keys of the JSON are:

- `genes`: This contains a list of genes in the index reference graph which are present and absent.
- `sample`: The value passed to the `-s/--sample` option.

- `susceptibility`: The keys of this entry are the drugs in the index catalogue. Each drug's entry contains `evidence` (Section S1.2) supporting the value in the `predict` (Section S1.1) section.

- `version`: the version of DrPRG and the index used.

#### S1.1 Predict

The `predict` entry for a drug is the resistance prediction for the sample. Possible values are:

- `S`: susceptible. This is the default prediction. If no mutations are detected for the sample, it is assumed to be susceptible.

- `F`: failed. Genotyping failed for one or more mutations for this drug. See the prediction VCF for more information.

- `U`: unknown. One or more mutations that are not present in the index catalogue were detected in a gene associated with this drug.

- `R`: resistant. One or more mutations from the index catalogue that confer resistance were detected.

- `u` or `r`: The same as the uppercase versions, but the mutation(s) were detected in a minor allele.

#### S1.2 Evidence

This is a list of the mutations supporting the prediction. The residue is one of `DNA` or `PROT` indicating whether the mutation describes a nucleotide or amino acid change, respectively. The variant is of the form `<ref><pos><alt>`; where `<ref>` is the reference sequence at position `<pos>` and `<alt>` is the nucleotide/amino acid the reference is changed to. See the catalogue documentation for more information. The `vcfid` is the value in the prediction VCF ID column for this mutation, making it easier to find a mutation in the VCF.

### S2 Sensitivity and specificity performance

**Table S1:** Comparison of resistance predictions from Illumina data for three tools. FN=number of false negatives, R=number of resistant isolates, FP=number of false positives, S=number of susceptible isolates, CI=confidence interval.

| Drug | Tool | FN(R) | FP(S) | Sensitivity (95% CI) | Specificity (95% CI) |
|---|---|---|---|---|---|
| Isoniazid | DrPRG | 1005(14671) | 586(25769) | 93.1% (92.7-93.5%) | 97.7% (97.5-97.9%) |
| | Mykrobe | 1076(14671) | 561(25769) | 92.7% (92.2-93.1%) | **97.8% (97.6-98.0%)** |
| | TBProfiler | 989(14671) | 649(25769) | **93.3% (92.8-93.7%)** | 97.5% (97.3-97.7%) |
| Rifampicin | DrPRG | 426(11776) | 620(28292) | 96.4% (96.0-96.7%) | 97.8% (97.6-98.0%) |
| | Mykrobe | 523(11776) | 604(28292) | 95.6% (95.2-95.9%) | **97.9% (97.7-98.0%)** |
| | TBProfiler | 370(11776) | 788(28292) | **96.9% (96.5-97.2%)** | 97.2% (97.0-97.4%) |
| Ethambutol | DrPRG | 595(6014) | 2290(27011) | 90.1% (89.3-90.8%) | 91.5% (91.2-91.8%) |
| | Mykrobe | 631(6014) | 2265(27011) | 89.5% (88.7-90.3%) | **91.6% (91.3-91.9%)** |
| | TBProfiler | 578(6014) | 2293(27011) | **90.4% (89.6-91.1%)** | 91.5% (91.2-91.8%) |
| Pyrazinamide | DrPRG | 776(3836) | 497(18050) | 79.8% (78.5-81.0%) | 97.2% (97.0-97.5%) |
| | Mykrobe | 798(3836) | 449(18050) | 79.2% (77.9-80.5%) | **97.5% (97.3-97.7%)** |
| | TBProfiler | 680(3836) | 513(18050) | **82.3% (81.0-83.4%)** | 97.2% (96.9-97.4%) |
| Levofloxacin | DrPRG | 268(3109) | 355(14867) | **91.4% (90.3-92.3%)** | 97.6% (97.4-97.8%) |
| | Mykrobe | 299(3109) | 330(14867) | 90.4% (89.3-91.4%) | **97.8% (97.5-98.0%)** |
| | TBProfiler | 276(3109) | 356(14867) | 91.1% (90.1-92.1%) | 97.6% (97.3-97.8%) |
| Moxifloxacin | DrPRG | 178(2260) | 1135(14698) | **92.1% (90.9-93.2%)** | 92.3% (91.8-92.7%) |
| | Mykrobe | 207(2260) | 1113(14698) | 90.8% (89.6-92.0%) | **92.4% (92.0-92.8%)** |
| | TBProfiler | 182(2260) | 1141(14698) | 91.9% (90.8-93.0%) | 92.2% (91.8-92.7%) |
| Ofloxacin | DrPRG | 138(781) | 69(6008) | **82.3% (79.5-84.8%)** | 98.9% (98.5-99.1%) |
| | Mykrobe | 147(781) | 62(6008) | 81.2% (78.3-83.8%) | **99.0% (98.7-99.2%)** |
| | TBProfiler | 138(781) | 65(6008) | **82.3% (79.5-84.8%)** | 98.9% (98.6-99.2%) |
| Amikacin | DrPRG | 269(1866) | 224(18737) | **85.6% (83.9-87.1%)** | 98.8% (98.6-99.0%) |
| | Mykrobe | 359(1866) | 192(18737) | 80.8% (78.9-82.5%) | **99.0% (98.8-99.1%)** |
| | TBProfiler | 270(1866) | 227(18737) | 85.5% (83.9-87.1%) | 98.8% (98.6-98.9%) |
| Capreomycin | DrPRG | 292(1300) | 299(13039) | **77.5% (75.2-79.7%)** | 97.7% (97.4-98.0%) |
| | Mykrobe | 367(1300) | 263(13039) | 71.8% (69.3-74.1%) | **98.0% (97.7-98.2%)** |
| | TBProfiler | 293(1300) | 305(13039) | 77.5% (75.1-79.7%) | 97.7% (97.4-97.9%) |
| Kanamycin | DrPRG | 361(2213) | 319(18382) | 83.7% (82.1-85.2%) | 98.3% (98.1-98.4%) |
| | Mykrobe | 445(2213) | 298(18382) | 79.9% (78.2-81.5%) | **98.4% (98.2-98.6%)** |
| | TBProfiler | 357(2213) | 322(18382) | **83.9% (82.3-85.3%)** | 98.2% (98.0-98.4%) |
| Streptomycin | DrPRG | 787(5365) | 681(10179) | 85.3% (84.4-86.3%) | 93.3% (92.8-93.8%) |
| | Mykrobe | 906(5365) | 676(10179) | 83.1% (82.1-84.1%) | 93.4% (92.9-93.8%) |
| | TBProfiler | 780(5365) | 661(10179) | **85.5% (84.5-86.4%)** | **93.5% (93.0-94.0%)** |
| Ethionamide | DrPRG | 733(2961) | 1023(11360) | **75.2% (73.7-76.8%)** | 91.0% (90.5-91.5%) |
| | Mykrobe | 847(2961) | 985(11360) | 71.4% (69.7-73.0%) | **91.3% (90.8-91.8%)** |
| | TBProfiler | 844(2961) | 995(11360) | 71.5% (69.8-73.1%) | 91.2% (90.7-91.7%) |
| Linezolid | DrPRG | 104(152) | 30(10915) | **31.6% (24.7-39.3%)** | 99.7% (99.6-99.8%) |
| | Mykrobe | 105(152) | 29(10915) | 30.9% (24.1-38.7%) | **99.7% (99.6-99.8%)** |
| | TBProfiler | 104(152) | 31(10915) | **31.6% (24.7-39.3%)** | 99.7% (99.6-99.8%) |
| Delamanid | DrPRG | 111(116) | 3(8154) | **4.3% (1.9-9.7%)** | 100.0% (99.9-100.0%) |
| | Mykrobe | 111(116) | 2(8154) | **4.3% (1.9-9.7%)** | **100.0% (99.9-100.0%)** |
| | TBProfiler | 111(116) | 2(8154) | **4.3% (1.9-9.7%)** | **100.0% (99.9-100.0%)** |

**Table S2:** Comparison of resistance predictions from Nanopore data for three programs. FN=number of false negatives, R=number of resistant isolates, FP=number of false positives, S=number of susceptible isolates, CI=confidence interval.

| Drug | Tool | FN(R) | FP(S) | Sensitivity (95% CI) | Specificity (95% CI) |
|---|---|---|---|---|---|
| Isoniazid | DrPRG | 9(60) | 5(48) | 85.0% (73.9-91.9%) | 89.6% (77.8-95.5%) |
| | Mykrobe | 9(60) | 4(48) | 85.0% (73.9-91.9%) | 91.7% (80.4-96.7%) |
| | TBProfiler | 9(60) | 3(48) | 85.0% (73.9-91.9%) | **93.8% (83.2-97.9%)** |
| Rifampicin | DrPRG | 5(57) | 1(44) | 91.2% (81.1-96.2%) | 97.7% (88.2-99.6%) |
| | Mykrobe | 5(57) | 1(44) | 91.2% (81.1-96.2%) | 97.7% (88.2-99.6%) |
| | TBProfiler | 5(57) | 1(44) | 91.2% (81.1-96.2%) | 97.7% (88.2-99.6%) |
| Ethambutol | DrPRG | 4(21) | 15(77) | **81.0% (60.0-92.3%)** | 80.5% (70.3-87.8%) |
| | Mykrobe | 4(21) | 15(77) | **81.0% (60.0-92.3%)** | 80.5% (70.3-87.8%) |
| | TBProfiler | 5(21) | 15(77) | 76.2% (54.9-89.4%) | 80.5% (70.3-87.8%) |
| Ofloxacin | DrPRG | 0(11) | 4(79) | 100.0% (74.1-100.0%) | 94.9% (87.7-98.0%) |
| | Mykrobe | 0(11) | 4(79) | 100.0% (74.1-100.0%) | 94.9% (87.7-98.0%) |
| | TBProfiler | 0(11) | 3(79) | 100.0% (74.1-100.0%) | **96.2% (89.4-98.7%)** |
| Amikacin | DrPRG | 0(14) | 3(81) | 100.0% (78.5-100.0%) | 96.3% (89.7-98.7%) |
| | Mykrobe | 0(14) | 3(81) | 100.0% (78.5-100.0%) | 96.3% (89.7-98.7%) |
| | TBProfiler | 0(14) | 3(81) | 100.0% (78.5-100.0%) | 96.3% (89.7-98.7%) |
| Capreomycin | DrPRG | 1(4) | 1(54) | 75.0% (30.1-95.4%) | 98.1% (90.2-99.7%) |
| | Mykrobe | 1(4) | 1(54) | 75.0% (30.1-95.4%) | 98.1% (90.2-99.7%) |
| | TBProfiler | 1(4) | 1(54) | 75.0% (30.1-95.4%) | 98.1% (90.2-99.7%) |
| Kanamycin | DrPRG | 0(3) | 1(55) | 100.0% (43.9-100.0%) | 98.2% (90.4-99.7%) |
| | Mykrobe | 0(3) | 1(55) | 100.0% (43.9-100.0%) | 98.2% (90.4-99.7%) |
| | TBProfiler | 0(3) | 1(55) | 100.0% (43.9-100.0%) | 98.2% (90.4-99.7%) |
| Streptomycin | DrPRG | 3(10) | 14(83) | 70.0% (39.7-89.2%) | 83.1% (73.7-89.7%) |
| | Mykrobe | 3(10) | 27(83) | 70.0% (39.7-89.2%) | 67.5% (56.8-76.6%) |
| | TBProfiler | 3(10) | 12(83) | 70.0% (39.7-89.2%) | **85.5% (76.4-91.5%)** |
| Ethionamide | DrPRG | 0(5) | 1(9) | 100.0% (56.6-100.0%) | 88.9% (56.5-98.0%) |
| | Mykrobe | 0(5) | 1(9) | 100.0% (56.6-100.0%) | 88.9% (56.5-98.0%) |
| | TBProfiler | 0(5) | 1(9) | 100.0% (56.6-100.0%) | 88.9% (56.5-98.0%) |
